# Gene-level, but not chromosome-wide, divergence between a very young house fly proto-Y chromosome and its homologous proto-X chromosome

**DOI:** 10.1101/2020.04.02.022707

**Authors:** Jae Hak Son, Richard P. Meisel

## Abstract

X and Y chromosomes are usually derived from a pair of homologous autosomes, which then diverge from each other over time. Although Y-specific features have been characterized in sex chromosomes of various ages, the earliest stages of Y chromosome evolution remain elusive. In particular, we do not know whether early stages of Y chromosome evolution consist of changes to individual genes or happen via chromosome-scale divergence from the X. To address this question, we quantified divergence between young proto-X and proto-Y chromosomes in the house fly, *Musca domestica*. We compared proto-sex chromosome sequence and gene expression between genotypic (XY) and sex-reversed (XX) males. We find evidence for sequence divergence between genes on the proto-X and proto-Y, including five genes with mitochondrial functions. There is also an excess of genes with divergent expression between the proto-X and proto-Y, but the number of genes is small. This suggests that individual proto-Y genes, but not the entire proto-Y chromosome, have diverged from the proto-X. We identified one gene, encoding an axonemal dynein assembly factor (which functions in sperm motility), that has higher expression in XY males than XX males because of a disproportionate contribution of the proto-Y allele to gene expression. The up-regulation of the proto-Y allele may be favored in males because of this gene’s function in spermatogenesis. The evolutionary divergence between proto-X and proto-Y copies of this gene, as well as the mitochondrial genes, is consistent with selection in males affecting the evolution of individual genes during early Y chromosome evolution.

## Introduction

In many organisms with two separate sexes, a gene on a sex chromosome determines whether an individual develops into a male or female. In XX/XY sex chromosome systems, males are the heterogametic sex (XY genotype), and females are homogametic with the XX genotype (Bull 1983). Most X and Y chromosomes are derived from a pair of ancestral autosomes. For example, one copy of the autosome can obtain a male-determining gene and become a proto-Y chromosome, and the homologous chromosome without the male-determiner becomes a proto-X. As the proto-X and proto-Y chromosomes diverge from each other over time, they become differentiated X and Y chromosomes (Bull 1983; Charlesworth et al. 2005). Sex chromosomes have originated and diverged from each other in multiple independent evolutionary lineages (Bachtrog et al. 2014; Beukeboom and Perrin 2014).

Despite their independent origins, non-homologous Y chromosomes share many common features across species (Charlesworth et al. 2005). First, “masculinization” occurs because male-limited inheritance of the Y chromosome favors the fixation of male-beneficial genetic variants (Rice 1996a). Second, suppressed recombination between the X and Y chromosomes evolves, possibly due to sexually antagonistic selection, meiotic drive, or genetic drift (Charlesworth 2017; Charlesworth 2018; Ponnikas et al. 2018). Third, “degeneration” occurs in nonrecombining regions—functional genes that were present on ancestral autosomes become pseudogenes on the Y chromosome because suppressed recombination between the X and Y inhibits the purging of deleterious mutations in Y-linked genes (Muller’s ratchet) and enhances the effects of hitchhiking (Charlesworth and Charlesworth 2000; Bachtrog 2013; Vicoso 2019). Other common features of Y chromosomes are repetitive sequences and enlarged heterochromatic regions due to reduced efficacy of purifying selection caused by suppressed recombination and a small effective population size (Skaletsky et al. 2003). In some cases, a mechanism evolves to compensate for the haploid dosage of X-linked genes in males, but this is not always the case (Mank 2013; Gu and Walters 2017).

Many features of Y chromosomes are thought to emerge shortly after an autosome obtains a new male-determining locus or becomes Y-linked. For example, recombination suppression has been considered to evolve after the emergence of a new sex-determining locus on a proto-Y chromosome to favor the co-inheritance of the sex-determining locus and male-beneficial/female-detrimental sexually antagonistic alleles (Orzack et al. 1980; van Doorn and Kirkpatrick 2007; Roberts et al. 2009; van Doorn and Kirkpatrick 2010). Additional sexually antagonistic alleles on the proto-Y chromosome are predicted to trigger progressive spread of the nonrecombining region along the chromosome (Rice 1987; van Doorn and Kirkpatrick 2007). Although these features have been characterized in sex chromosomes of various ages and degeneration levels (Bachtrog 2013; Zhou et al. 2014), the very first stages of Y chromosome evolution are poorly understood because of a lack of extremely young sex chromosome systems. Recent studies of young sex chromosomes have identified multiple types of X-Y differentiation, including suppressed recombination, Y chromosome gene loss, and X chromosome dosage compensation (Bergero et al. 2013; Mahajan et al. 2018; Darolti et al. 2019; Krasovec et al. 2019), which makes it challenging to determine which type of differentiation occurs first.

The extent to which the early evolution of sex chromosomes is dominated by chromosome-wide X-Y divergence versus changes in individual genes remains unclear. This study addresses that shortcoming by determining how a young proto-Y chromosome has differentiated from its homologous proto-X chromosome shorty after its emergence. We are especially interested in how gene expression differences accumulate between the proto-Y and proto-X chromosomes. As the proto-Y and proto-X chromosomes diverge, it is expected that alleles on the proto-Y chromosome are up-or down-regulated because of *cis*-regulatory sequence differences that contribute to proto-Y gene expression (Zhou and Bachtrog 2012a; Zhou and Bachtrog 2012b; Wei and Bachtrog 2019). These *cis*-regulatory effects may be especially important for the expression of sexually antagonistic (male-beneficial/female-deleterious) alleles and degeneration of functional genes (Rice 1984; Zhou and Bachtrog 2012a). Degeneration of Y-linked genes has been shown to be accompanied by decreased expression as a result of relaxed selective constraints (Zhou and Bachtrog 2012a; Wei and Bachtrog 2019). However, the accumulation of gene expression differences separately from degeneration during the very earliest stages of sex chromosome evolution are not well understood.

We used the house fly, *Musca domestica*, as a model system to study the early evolution of sex chromosomes because it has very young proto-sex chromosomes that are still segregating as polymorphisms within natural populations (Hamm et al. 2015). The *M. domestica male determiner* (*Mdmd*) can be found on what was historically called the Y chromosome (Y^M^) and on at least three other chromosomes (Sharma et al. 2017). The house fly Y^M^ (and X) chromosome has fewer than 100 genes, while the other five chromosomes each have >2,000 genes (Meisel and Scott 2018). Each chromosome carrying *Mdmd*, including Y^M^, is a recently derived proto-Y chromosome (Meisel et al. 2017). *Mdmd* arose in the house fly genome after divergence from stable fly (*Stomoxys calcitrans*) and horn fly (*Haematobia irritans*), within the past 27 million years (Sharma et al. 2017; Meisel et al. 2020). This provides an upper-bound on the age of the house fly proto-Y chromosomes, although the minimal sequence and morphological divergence between the proto-Y and proto-X chromosomes (Boyes et al. 1964; Hediger et al. 1998; Meisel et al. 2017) suggest they are much younger than that. It is not clear the extent to which the house fly proto-Y chromosomes are masculinized or degenerated. A previous study revealed a small, but significant, effect of the proto-Y chromosomes on gene expression (Son et al. 2019). However, it could not resolve if the expression differences are the result of changes in the expression of the proto-Y copies, proto-X copies, or both.

In this study, we tested if one house fly proto-Y chromosome, the third chromosome carrying *Mdmd* (III^M^), has evidence of gene-by-gene or chromosome-wide differentiation from its homologous proto-X chromosome by evaluating DNA sequence and gene expression differences between proto-Y genes and their proto-X counterparts. We selected this proto-sex chromosome (as opposed to other house fly proto-Y chromosomes) for three reasons. First, III^M^ is one of the two most common proto-Y chromosomes found in natural populations (Hamm et al. 2015). Second, the other common proto-Y (known as Y^M^) has fewer than 100 genes, which is over one order of magnitude less than III^M^ (Meisel et al. 2017; Meisel and Scott 2018). More genes on the third chromosome gives us greater power to detect divergence between the proto-Y and proto-X. Third, we are able to create sex-reversed males (genotypic females that are phenotypically male) with the same genetic background as III^M^ males (Hediger et al. 2010). This allows us to compare gene expression between phenotypic males that only differ in whether or not they carry the III^M^ proto-Y chromosome (Son et al. 2019).

Our expectation for the extent of gene-by-gene versus chromosome-wide differentiation between the proto-X and proto-Y chromosomes will depend on the extent of X-Y recombination. We expect gene-by-gene differentiation on young sex chromosomes if recombination prevents the effects of Muller’s ratchet and genetic hitchhiking (Charlesworth and Charlesworth 2000). Recombination between the house fly proto-X and proto-Y is possible because, unlike *Drosophila*, there may be recombination in male house flies (Feldmeyer et al. 2010). In addition, female house flies can carry a proto-Y chromosome if they have a female-determining allele on another chromosome (Mcdonald et al. 1978; Hediger et al. 2010; Hamm et al. 2015), which would provide additional opportunities for recombination. However, if there are recombination suppressors (such as chromosomal inversions that differentiate the proto-Y and proto-X), this would promote chromosome-wide divergence between the proto-X and proto-Y via Muller’s ratchet and hitchhiking (Ponnikas et al. 2018).

## Results and Discussion

### DNA sequence divergence between the proto-Y and proto-X chromosomes

We used RNA-seq data to identify single nucleotide polymorphisms (SNPs) and small insertions/deletions (indels) within genes in genotypic (III^M^/III) and sex-reversed (III/III) male house flies (Son et al. 2019). For each gene, we counted the number of sites that are heterozygous in either genotypic or sex-reversed males (i.e., two alleles in at least one genotype). We then calculated the percent of those sites (per gene) that are heterozygous only in the genotypic males. This value is 0% if heterozygous sites are only in sex-reversed males, it is 100% if heterozygous sites are only in genotypic males, and it is 50% if the same number of heterozygous sites are found in both genotypes. We found that the genotypic males have an excess of heterozygous sites in third chromosome genes, relative to the sex-reversed males (Figure 1; *P* < 10^−16^ in a Wilcoxon rank sum test comparing percent heterozygous sites in genes on the third chromosome with genes on the other chromosomes). This is consistent with elevated third chromosome heterozygosity in a previous comparison between III^M^ males and Y^M^ males (Meisel et al. 2017), and it suggests that the sequences of genes on the III^M^ proto-Y chromosome are differentiated from the copies on the proto-X (i.e., the standard third chromosome).

**Figure 1.**
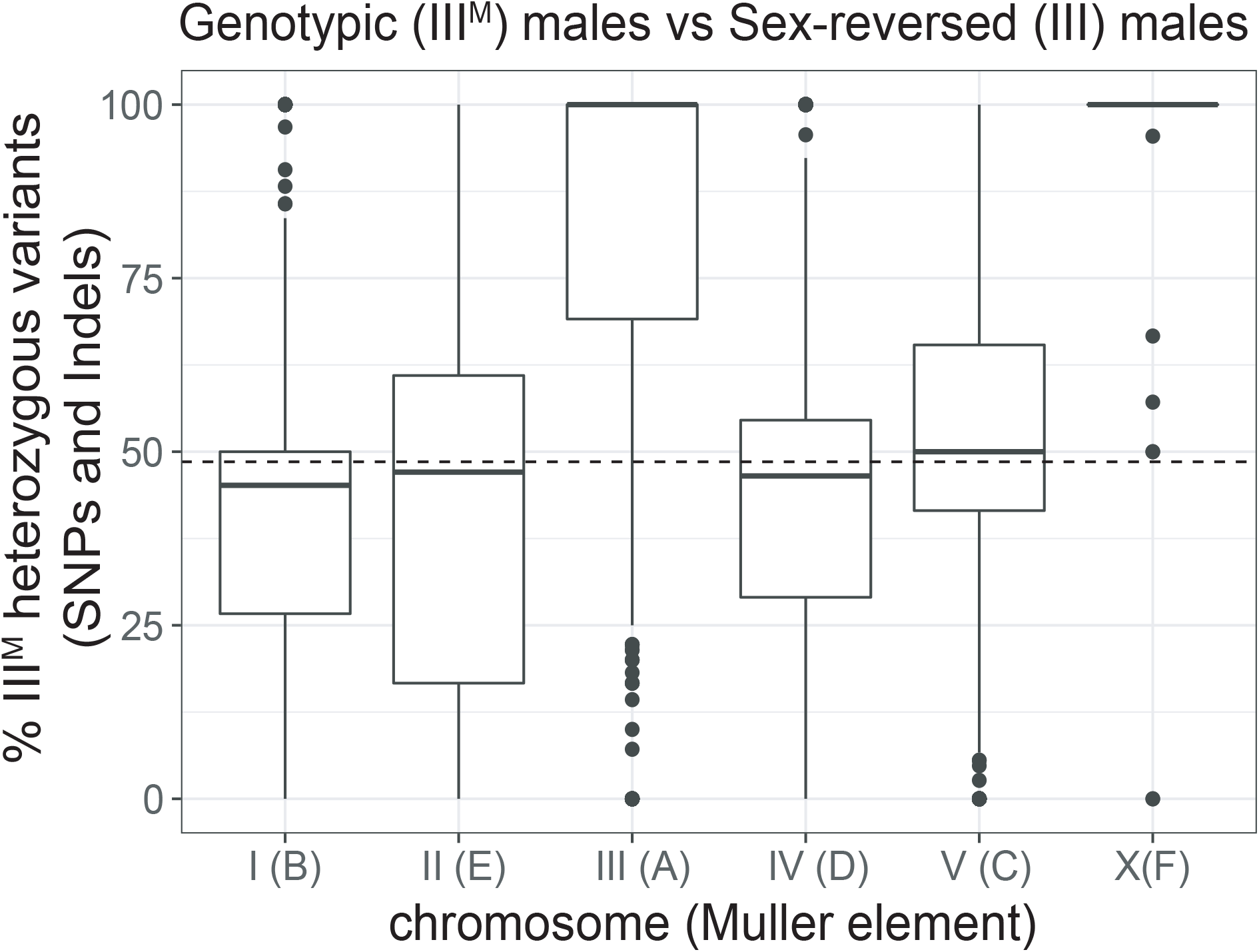
Elevated heterozygosity on the third and X chromosomes in genotypic (III^M^/III) males relative to sex-reversed (III/III) males. The boxplots show the distributions of the percent of heterozygous variants per gene in the genotypic males relative to the sex-reversed males (% III^M^ heterozygous variants) on each chromosome. Muller element nomenclature for each chromosome is shown in parentheses (Meisel and Scott 2018). See Materials and Methods for the calculation of % III^M^ heterozygous variants. Values more than 50% indicate more heterozygous variants in genotypic (III^M^/III) males, and less than 50% indicates more heterozygous variants in sex-reversed males. The median across autosomes I, II, IV, and V is represented by a dashed line.

We expect that the ancestral X chromosome would have the same levels of heterozygosity in genotypic males (X/X; III^M^/III) and sex-reversed males (X/X; III/III) due to the presence of two copies of the X chromosome in both genotypes. However, the III^M^ males have elevated heterozygosity on the X chromosome (Figure 1; *P* = 8.32×10^−13^ in a Wilcoxon rank sum test comparing the X chromosome with chromosomes I, II, IV, and V). Elevated X chromosome heterozygosity in III^M^ males was also observed in a comparison with Y^M^ males (Meisel et al. 2017), and its cause remains unresolved. One possible explanation for elevated X chromosome heterozygosity is that the III^M^ chromosome was created by a fusion between the third chromosome and the Y^M^ chromosome. Because Y^M^ has nearly identical gene content as the X chromosome (Meisel et al. 2017), a III-Y^M^ fusion would cause III^M^ males to have three copies of X chromosome genes, increasing the likelihood that III^M^ males are heterozygous at any given site on the X chromosome. This hypothesis remains to be tested.

We further tested for divergence between the III^M^ proto-Y chromosome and its homologous proto-X by assembling a III^M^ male genome using Oxford Nanopore reads from the same strain as our RNA-seq data. We were specifically interested in identifying contigs that were separately assembled from homologous regions on the proto-X and proto-Y chromosomes (i.e., gametologs; Garcia-Moreno and Mindell 2000). Assembly into separate contigs would provide evidence for divergence between the proto-X and proto-Y sequences. Our III^M^ male assembly is smaller (427 Mb) than the reference house fly genome assembly (691 Mb; Scott et al. 2014) and flow cytometry estimates of genome size (~1,000 Mb; Picard et al. 2012), suggesting that we are missing 30-50% of the genome sequence in our assembly. The reduced assembled size of our III^M^ male genome is likely the result of low sequencing coverage. Nonetheless, we should be able to identify X-Y divergence, albeit with reduced power relative to a more complete genome assembly.

We took two approaches to identify separate proto-X and proto-Y contigs in our assembly: 1) identifying genes on the same contig as the male-determining gene (*Mdmd*); and 2) testing for contigs containing sequences that are enriched in males relative to females (Carvalho and Clark 2013). Our approaches will identify regions on the proto-sex chromosome that contain genes that can be assigned to the third chromosome, are present on both the proto-X and proto-Y with sufficient divergence to assemble separately, and with both the proto-X and proto-Y gametologs assembled in our III^M^ male genome. We identified two loci, one from each of our two approaches (see Supplementary Materials for details), with sufficient X-Y divergence to assemble into separate proto-X and proto-Y contigs (Figure 2). At each locus, we have one proto-Y contig and one proto-X contig.

**Figure 2.**
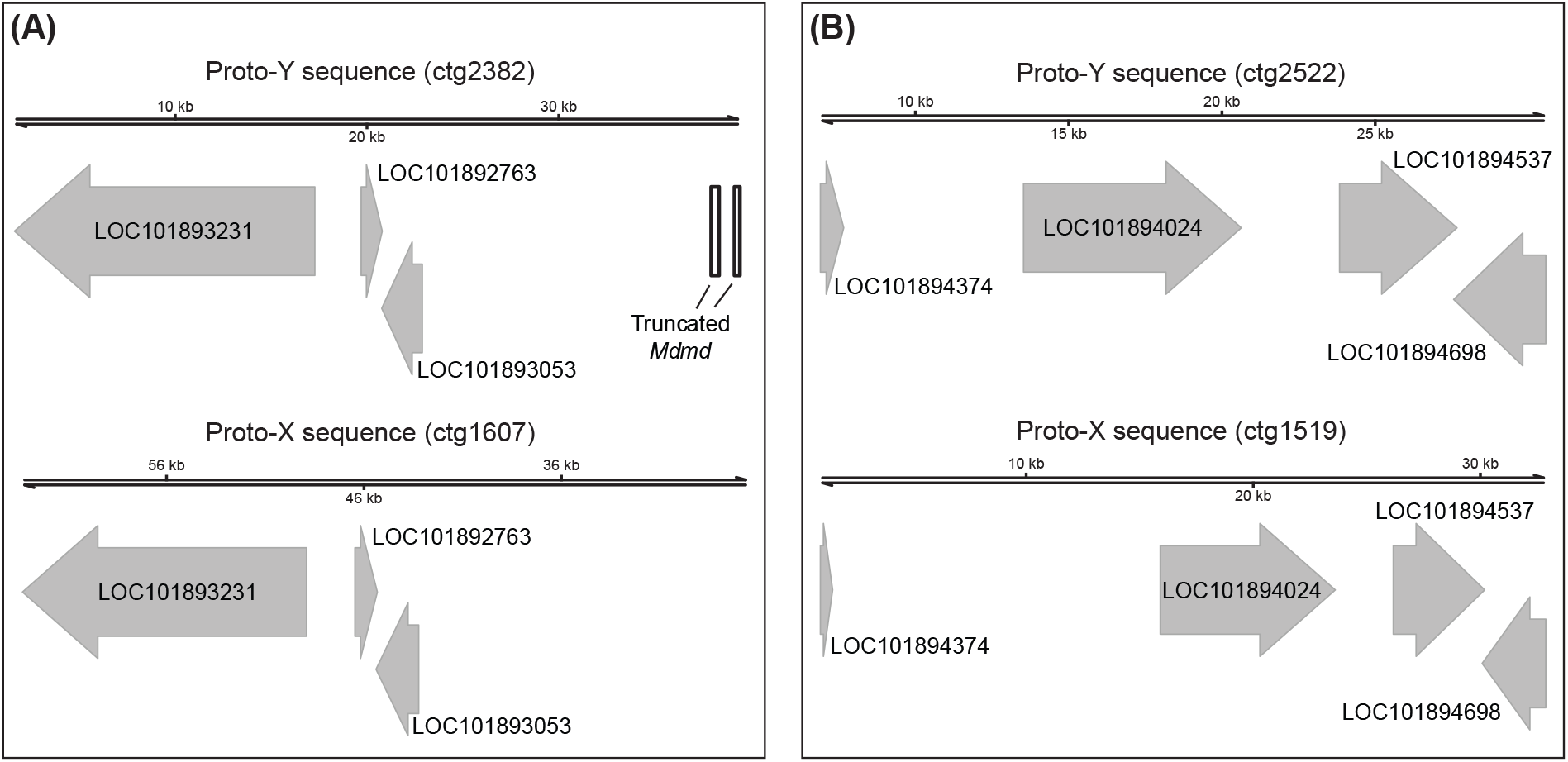
Two proto-X and proto-Y loci identified in the III^M^ genome assembly. (A) One locus was identified with a truncated *Mdmd* on the proto-Y (ctg2382) and the same three genes on both the proto-X (ctg1607) and proto-Y. (B) One locus was identified with four genes on both the proto-X (ctg1519) and proto-Y (ctg2522). The two contigs (ctg2522 and ctg1519) were assigned to the proto-Y and proto-X based on sequences that are enriched in the male relative to female reads (see Supplementary Figure 1).

One of the three genes on one proto-Y contig (ctg2382) and all four genes on the other proto-Y contig (ctg2522) are nuclear-encoded mitochondrial genes (Figure 2 and Supplementary Table 2). Nuclear-encoded mitochondrial genes can evolve under sexually antagonistic selection (Rand et al. 2001), possibly because of conflicts over mitochondrial functions in sperm and in other tissues (Gemmell et al. 2004; Gallach et al. 2010). These inter-sexual conflicts could favor or select against X-linked mitochondrial genes depending on the extent of co-transmission of the X chromosome and mitochondria in females (Drown et al. 2012; Dean et al. 2014). Our results suggest that mito-nuclear inter-sexual conflicts might also be important for X-Y divergence in young sex chromosomes, where sexually antagonistic variants could be partitioned between female-beneficial X-linked alleles and male-beneficial Y-linked alleles. Additional work is required to test this hypothesis.

Two of the genes on the proto-Y contigs have differences in their protein-coding sequence from their proto-X chromosome gametologs. One of the two (*LOC101894698*, encoding fast kinase domain-containing protein 5, mitochondrial) contains three missense variable sites at which genotypic (III^M^/III) males are heterozygous and sex-reversed (III/III) male are homozygous (Supplementary Table 3). The protein encoded by this gene contains an RNA-binding (RAP) domain, but all three missense variants are not found in the domain. The other gene (*LOC101893231*, which does not have a predicted mitochondrial function) has three missense alleles at which genotypic males are heterozygous and sex-reversed male are homozygous (Supplementary Table 3). One of three missense sites (at position 29,021 in the scaffold of the reference genome) in this gene is found in a proline aminopeptidase P II domain. For all missense alleles in both genes, we inferred the III allele as the one in common between genotypic and sex-reversed males and the III^M^ allele as the one unique to genotypic males. However, the inferred III^M^ alleles are not specific to the III^M^ chromosome because all III^M^ alleles in *LOC101894698* and one of the III^M^ alleles in *LOC101893231* (at position 29,021) were found in the reference genome (which comes from a genotypic female without a III^M^ chromosome). Therefore, some of the III^M^ alleles in these genes are segregating as polymorphic variants on the proto-X chromosome.

An alternative explanation of our candidate proto-Y contigs (ctg2382 and ctg2522) is that they are paralogous sequences that have been duplicated on the proto-Y chromosome, creating a second Y-linked copy of the genes within the duplicated region. Intra-chromosomal duplications are a common feature of Y chromosomes (Skaletsky et al. 2003; Hughes et al. 2010; Soh et al. 2014; Bachtrog et al. 2019; Ellison and Bachtrog 2019). It is therefore possible that the sequences we classify as on the proto-X are actually present on both the proto-X and proto-Y, while the contigs we classify as proto-Y sequences are intra-chromosomal duplications on the proto-Y. However, even in this scenario, the proto-Y sequences are still unique to the proto-Y. Therefore, our interpretations are unlikely to be affected by whether the sequences we classify as proto-Y are true gametologs of the proto-X or intra-chromosomal duplications on the proto-Y.

### Gene expression divergence between the proto-Y and proto-X chromosomes

We next tested if differences in *cis*-regulatory sequences between the proto-Y and proto-X chromosomes contribute to expression differentiation. We quantified differential expression between the proto-X and proto-Y chromosome copies of the third chromosome genes by measuring allele-specific expression (ASE) in normal (genotypic) males carrying a III^M^ proto-Y chromosome and sex-reversed (III/III) males with no proto-Y chromosome. Comparing genotypic and sex-reversed males allows us to control for the effect of sexually dimorphic gene expression on the inference of divergence between the proto-Y (III^M^) and proto-X (III) chromosomes.

To quantify ASE of genes in genotypic (III^M^/III) and sex-reversed (III/III) males, we used existing RNA-seq data (Son et al. 2019) along with our new Oxford Nanopore long read sequencing data. We used the IDP-ASE pipeline (Deonovic et al. 2017), which is more accurate when a hybrid of short and long reads is provided as input. This is because the long reads are used to construct haplotypes, which are used to estimate ASE from the RNA-seq data (see Materials and Methods). This approach differs from our comparison of proto-X and proto-Y contigs because the ASE analysis does not require separate assembly of proto-X and proto-Y contigs. Instead, IDP-ASE uses the raw long read sequences from genotypic and sex reversed males to phase haplotypes when inferring ASE. We measured ASE as the proportion of iterations in a Markov chain Monte Carlo (MCMC) simulation in which the expression of a focal haplotype is estimated as >0.5. This proportion gives a measure of ASE ranging from 0 (extreme ASE in favor of one allele) to 1 (extreme ASE in favor of another allele), with 0.5 indicating equal expression of both alleles. We are unable to determine if the focal haplotype refers to the III^M^ or III allele across the entire third chromosome, but we do differentiate between these alleles for a handful of genes where we are able to perform manual curation (see below).

We assigned each gene with sufficient expression data into one of five bins of ASE. The proportions of iterations with focal haplotypes >0.5 were overrepresented at five values (0, 0.25, 0.5, 0.75, and 1) both in genotypic and sex-reversed males (Supplementary Figure 2 and 3). These proportions may be overrepresented because we only sampled two genotypes for our ASE analysis, which caused us to have a non-continuous distribution of proportions. We divided the proportion of iterations with the focal haplotypes >0.5 into five bins, with each bin capturing one of the five most common proportions (Supplementary Figure 2 and 3): 1) extreme ASE, with a value between 0 and 0.125; 2) moderate ASE, with a value between 0.125 and 0.375; 3) non-ASE, with a value between 0.375 and 0.625; 4) moderate ASE, with a value between 0.625 and 0.875; and 5) extreme ASE, with a value between 0.875 and 1. In the analysis below, we considered a gene to have ASE if it falls into one of the two bins of extreme ASE; genes in the non-ASE bin (proportion of focal haplotype >0.5 between 0.375 and 0.625) were classified as having non-ASE. Genes with moderate ASE were excluded from most of our analyses in order to be conservative about ASE assignment.

We first investigated ASE of genes we identified at the two loci where we have separate proto-X and proto-Y contigs from our genome assembly (Supplementary Table 2). We had enough sequence coverage and heterozygous sites to estimate ASE in 5/7 genes. Two mitochondrial genes (*LOC101894537* and *LOC101894698*) had extreme ASE in genotypic males and moderate ASE in sex-reversed males. The elevated ASE in genotypic males is suggestive of expression divergence between the proto-Y and proto-X copies. Three genes (*LOC101892763*, *LOC101893231*, and *LOC101894024*) exhibited extreme ASE in sex-reversed males and non-ASE or moderate ASE in genotypic males. Elevated ASE in sex-reversed males is not expected because they are genotypic females with two copies of the proto-X chromosome. This result suggests there is not large-scale expression divergence between the proto-Y and proto-X chromosomes. The limited expression divergence (Supplementary Table 2) and paucity of fixed amino acid differences (Supplementary Table 3) between proto-Y and proto-X gametologs are consistent with minimal differentiation between the house fly proto-Y and proto-X chromosomes.

We next examined ASE across the entire third chromosome. If the III^M^ proto-Y chromosome is differentiated in gene expression from its homologous III proto-X chromosome because of differences in *cis-*regulatory alleles across the entire third chromosome, then we expect a higher fraction of genes with ASE on the third chromosome in the genotypic (III^M^/III) males than in the sex-reversed (III/III) males. In contrast to that expectation, we did not find an excess of genes with ASE in genotypic males compared to ASE genes in sex-reversed males on the third chromosome relative to other chromosomes (Figure 3A and Supplementary Table 4; Fisher’s exact test, *P* = 0.6996). This result suggests that the III^M^ proto-Y chromosome is not broadly differentiated in *cis*-regulatory alleles from the standard third (proto-X) chromosome. This provides evidence that the early stages of Y chromosome evolution do not involve chromosome-wide changes in gene regulation.

**Figure 3.**
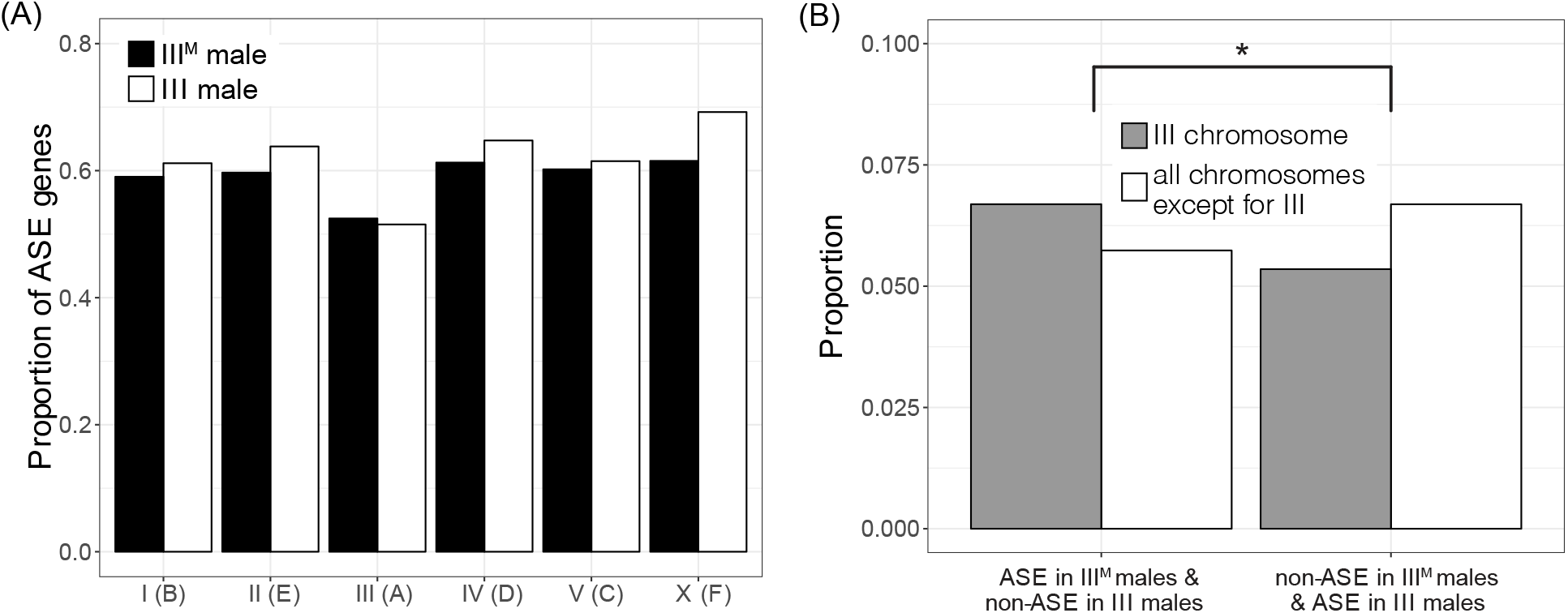
Evidence for moderately elevated ASE on the third (proto-Y) chromosome in III^M^ males. (A) Proportions of genes with ASE in genotypic (III^M^) or sex-reversed (III) males on each chromosome. There is not a significant difference on any chromosome between the two genotypes. (B) Proportions of genes with ASE in genotypic males and non-ASE in sex-reversed males on the third chromosome and all other chromosomes (left two bars). Proportions of genes with non-ASE in genotypic males and ASE in sex-reversed males on the third chromosome and all other chromosomes (right two bars). The asterisk indicates a significant difference (*p*<0.05) in the number of genes in these categories as determined by Fisher’s exact test.

We next identified individual genes with differences in ASE between genotypic (III^M^/III) and sex-reversed (III/III) males. There are 95 third chromosome genes with ASE in the genotypic males that are non-ASE in the sex-reversed males (Supplementary Table 5). These genes could have ASE in III^M^ males because of differences in *cis* regulatory sequences between the III^M^ and standard third chromosome. To test whether the observed number of third chromosome genes with ASE in genotypic males that are non-ASE in sex-reversed males is in excess of a null expectation, we determined the number of third chromosome genes with ASE in sex-reversed males that are non-ASE in genotypic males (i.e., the opposite of what we did above to find the first set of 95 genes). There are 76 third chromosome genes with ASE in the sex-reversed males that are non-ASE in genotypic males (Supplementary Table 5). We also identified 241 genes on other chromosomes with ASE in genotypic males that are non-ASE in sex-reversed males, as well as 281 genes on other chromosomes with ASE in sex-reversed males that are non-ASE in genotypic males (Figure 3B and Supplementary Table 5). We do not expect any difference in ASE between genotypic and sex reversed males for chromosomes other than the third. Comparing genes with and without ASE on the third chromosome and the rest of the genome, there is indeed an excess of third chromosome genes with ASE in genotypic males that are non-ASE in sex-reversed males (Figure 3B and Supplementary Table 5; Fisher’s exact test, *P* = 0.03467). These results suggest that, while the III^M^ proto-Y chromosome is not broadly differentiated in *cis*-regulatory sequences from the standard third (proto-X) chromosome, there is an excess of individual genes with *cis*-regulatory differences between the III^M^ proto-Y and its homologous proto-X chromosome.

Male-specific selection on individual proto-Y genes could be responsible for ASE in genotypic III^M^ males. In this scenario, male-specific selection would favor *cis-*regulatory variants that drive up-or down-regulation of the III^M^ copy of a gene (Parsch and Ellegren 2013; Mank 2017). These sex-specific selection pressures are expected to have the greatest effect for genes closest to the male-determining *Mdmd* locus because those genes are most likely to be co-inherited with *Mdmd* (Charlesworth et al. 2014). Unfortunately, we lack a chromosome-scale assembly of the house fly genome (Scott et al. 2014; Meisel and Scott 2018), which prevents us from testing if genes with ASE are clustered on the third chromosome in close proximity to the *Mdmd* locus.

### Up-regulation of the Y-allele and male-biased expression of a testis-expressed gene

We next tested if genes with ASE are differentially expressed between genotypic males, sex-reversed males, and females. A relationship between ASE and sexually dimorphic expression would suggest that up-or down-regulation of Y-linked alleles affects sexually dimorphic phenotypes. We started by selecting genes that we had previously identified as differentially expressed between genotypic (III^M^/III) and sex-reversed (III/III) males (Son et al. 2019). These two genotypes have nearly the same expression profiles, with a small number of differentially expressed genes. We were specifically interested in genes on the third (proto-sex) chromosome with “discordant sex-biased expression”. Discordant sex-biased genes have male-biased expression in genotypic males (i.e., upregulated relative to phenotypic females) and female-biased expression in the sex-reversed males (downregulated relative to phenotypic females), or vice versa. This pattern of expression is suggestive of *cis*-regulatory divergence between the proto-Y and proto-X chromosomes—a hypothesis that was presented but not tested in our previous study (Son et al. 2019). Here, we test that hypothesis by determining if any third chromosome genes with discordant sex-biased expression have ASE consistent with *cis-* regulatory divergence between the proto-Y (III^M^) and proto-X (III) alleles.

We identified a single gene (*LOC101899975*) with discordant sex-biased gene expression out of the 95 genes on the third chromosome with ASE in the genotypic (III^M^/III) males that are non-ASE in the sex-reversed (III/III) males. This gene is homologous to *dynein assembly factor 5, axonemal* (human gene *DNAAF5* and *Drosophila melanogaster* gene *HEATR2*). The gene, which we refer to as *M. domestica HEATR2* (*Md-HEATR2*), is expected to encode a protein that functions in flagellated sperm motility (Diggle et al. 2014), and it has strong testis-biased expression in *D. melanogaster* (Chintapalli et al. 2007). *Md-HEATR2* has male-biased expression in the abdomens of genotypic males and female-biased expression in the abdomens of sex-reversed males (Son et al. 2019), suggesting that expression differences between the III^M^ proto-Y and the standard third (proto-X) chromosome cause the male-biased expression of the gene in the genotypic males.

We identified three diagnostic variant sites for ASE within *Md-HEATR2* (Figure 4A), which are all synonymous SNPs. The genotypic (III^M^/III) males are heterozygous and the sex-reversed (III/III) males are homozygous at all diagnostic sites. We inferred the allele on the standard third chromosome as the one in common between genotypic and sex-reversed males, and the III^M^ allele as the one unique to genotypic males at each diagnostic variant site. Curiously, all three III^M^ alleles are found in the reference genome, suggesting that these synonymous variants are not fixed differences between the proto-Y and proto-X. *Md-HEATR2* is expressed higher in III^M^ genotypic males than in sex-reversed males (Figure 4A). In the III^M^ genotypic males, the III^M^(Y-linked) alleles are expressed higher than the X-linked alleles, indicating that the Y-linked alleles are associated with the up-regulation of the gene in III^M^ genotypic males relative to sex-reversed males (Figure 4A). The copy of *Md-HEATR2* on the III^M^ proto-Y chromosome is therefore up-regulated relative to the proto-X copy, consistent with higher expression of *Md-HEATR2* in genotypic males.

**Figure 4.**
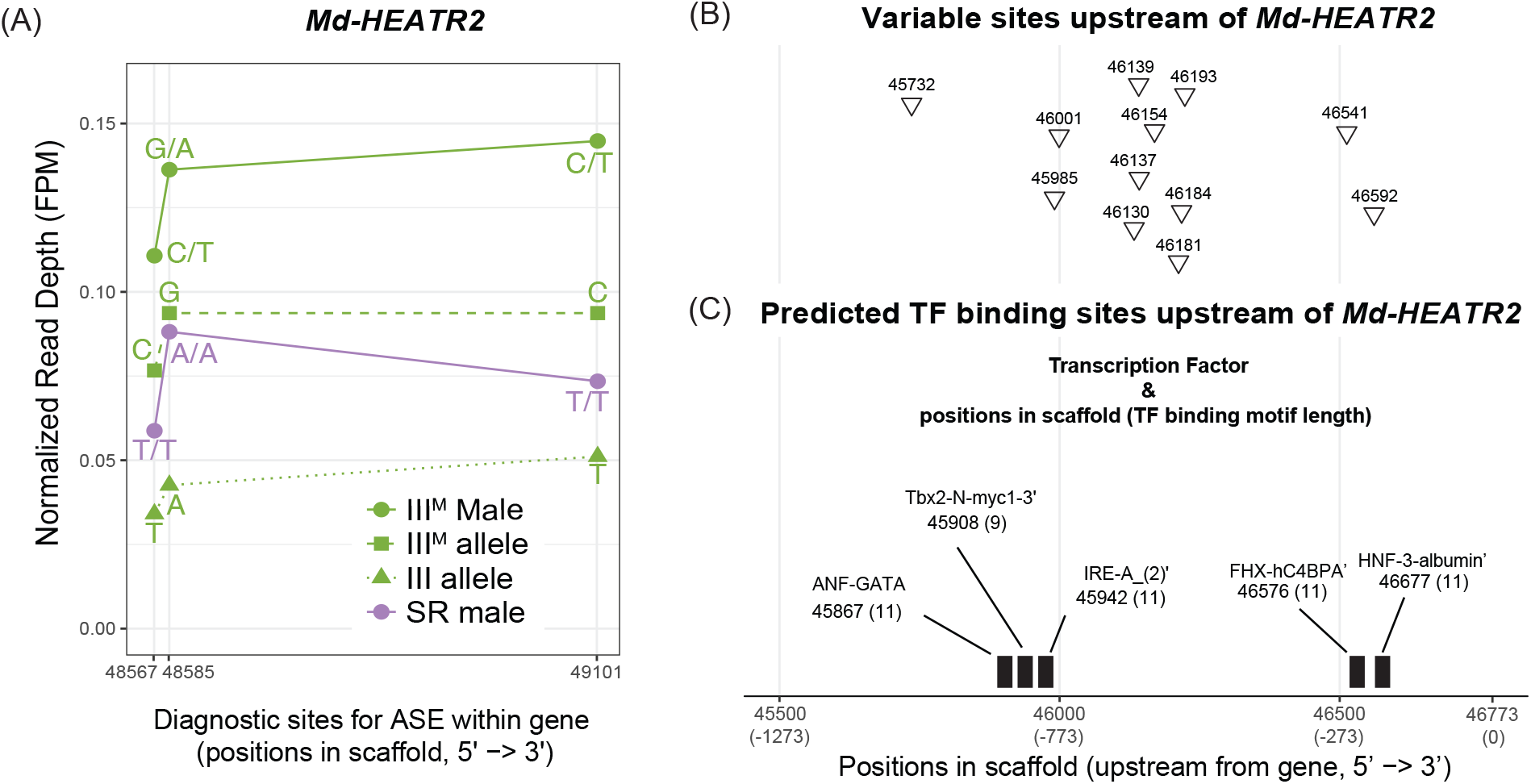
Allele-specific expression (ASE) of *Md-HEATR2*. (A) Diagnostic variable sites for ASE in the *Md-HEATR2* gene are based on haplotypes estimated in IDP-ASE. Read depth is measured as fragments per million mapped reads (FPM) in III^M^ males, sex-reversed (SR) males, and the III^M^ or III allele in III^M^ males. (B) Variable sites that differ between genotypic (III^M^) males and sex-reversed males (triangles) across 1,273 base pairs upstream of *Md-HEATR2* were identified using Oxford Nanopore long reads only. (C) Transcription factor (TF) binding motifs predicted within 1,273 base pairs upstream of *Md-HEATR2*. The starting position of each motif is given, along with its length (in parentheses). None of the variable sites overlap with predicted TF motifs. All positions are coordinates in scaffold NW_004764965.1 of the reference genome assembly (Scott et al. 2014).

Using our Nanopore sequencing reads mapped to the reference genome, we examined 1,273 base pairs upstream of *Md-HEATR2* to identify diagnostic sites that could be responsible for regulating the expression differences between the proto-X and proto-Y alleles. We chose that distance because it includes the first variable site we could identify on the scaffold containing *Md-HEATR2* in our Nanopore data (i.e., including a larger region would not provide any additional information). We found twelve variable sites with different alleles (SNPs and small indels) between genotypic (III^M^/III) and sex-reversed (III/III) males (Figure 4B). We next examined whether these sites are located within a potential transcription factor (TF) binding region. We found five TF binding regions predicted upstream of *Md-HEATR2* using the ‘Tfsitescan’ tool in the ‘object-oriented Transcription Factors Database’ (Ghosh 2000). However, none of the twelve variable sites are found within any predicted TF binding regions (Figure 4C). It is possible that the *cis*-regulatory sequences responsible for differential expression are located outside of the region we were able to investigate, which would be consistent with our failure to identify fixed differences in the exons between the proto-Y and proto-X copies of *Md-HEATR2*. A long distance would reduce the genetic linkage between the *cis*-regulatory region and transcribed sequence, allowing for the X-Y differentiation of the regulatory region without differentiation of the transcribed gene. Further work is needed to determine how the differential expression of the proto-X and proto-Y copies of *Md-HEATR2* is regulated.

We hypothesize that the upregulation of the proto-Y copy of *Md-HEATR2* is the result of selection for higher expression in males. We think that the *cis-*regulatory region is under selection, rather than the protein-coding sequence, because we fail to find any protein coding differences between the proto-X and proto-Y copies, and the synonymous differences we observe are not fixed differences. HEATR2 is involved in dynein arm assembly (Diggle et al. 2014), and axonemal dynein is essential for flagellated sperm motility (Kurek et al. 1998; Carvalho et al. 2000). Therefore, it may be beneficial to male fitness to have higher expression of *Md-HEATR2* in testis. However, HEATR2 also functions in mechanosensory neurons in *Drosophila* (Diggle et al. 2014). There may be conflict over the *cis*-regulatory sequences that promote expression in testis and neurons, which may prevent up-regulation of the proto-X copy of *Md-HEATR2* in testis. These opposing (i.e., sexually antagonistic) selection pressures on *Md-HEATR2* expression would be resolved in a Y-linked copy that is only under selection in males (Rice 1996b). Alternatively, the expression differences between sex-reversed and genotypic males could be caused by differences in tissue scaling between genotypes (Montgomery and Mank 2016). For example, if genotypic males have larger testes and if the proto-Y allele is preferentially expressed in testis, then it will appear as if genotypic males have higher expression of *Md-HEATR2*. Even in this model, there is ASE associated with testis-specific expression, which is consistent with male-specific selection favoring the proto-Y allele.

The up-regulation or testis-biased expression of the proto-Y copy of *Md-HEATR2* may not completely resolve sexually antagonistic selection pressures on *Md-HEATR2* expression because of three notable features of house fly genetics. First, Y-linked alleles are only under male-specific selection when they are in complete genetic linkage with a male-determining gene (Charlesworth et al. 2014). Complete linkage between loci can be caused by recombination suppressors (e.g., chromosomal inversions) that prevent genetic exchange between the X and Y chromosomes (Rice 1987; Charlesworth 2017). While there is no direct evidence for inversions or other suppressors of recombination on the house fly third chromosome, crossing over in male meiosis is rare in most flies (Gethmann 1988). A general lack of recombination in males would prevent X-Y exchange. However, male recombination has been documented in house fly (Feldmeyer et al. 2010), suggesting that X-Y exchange is possible. Second, female house flies can carry proto-Y chromosomes if they also carry the female-determining *Md-tra^D^* allele (Hamm et al. 2015). Female-transmission of the proto-Y provides another avenue for X-Y exchange in the absence of an inversion or other suppressor of recombination on the proto-Y or proto-X. We observe alleles in *Md-HEATR2* and other third chromosome genes that are found on both the proto-Y and proto-X (Supplementary Table 3), which is consistent with either X-Y recombination or ancestral polymorphisms that predate the formation of the proto-sex chromosome. Third, when females carry a proto-Y chromosome, there will be selection against male-beneficial, female-detrimental sexually antagonistic alleles, which will further prevent the resolution of sexual conflict.

Selection on proto-Y alleles in females, along with X-Y recombination that moves proto-X alleles onto the proto-Y, will reduce the efficacy of male-specific selection pressures on sexually antagonistic alleles of *Md-HEATR2*. The extent of these effects will depend on the frequency of proto-Y chromosomes found in females. First, the extent to which selection in females acts against male-beneficial sexually antagonistic alleles depends on the frequency with which the proto-Y chromosomes are found in females (Rice 1984). Second, if recombination rate is sexually dimorphic, the rate of population level recombination between the male-determiner and *Md-HEATR2* will depend on how often the proto-Y chromosomes are found in females (Sardell et al. 2018). Notably, the frequencies of proto-Y chromosomes and the female-determining *Md-tra^D^* allele vary across populations (Hamm et al. 2015; Meisel et al. 2016). Therefore, the frequency with which females carry proto-Y chromosomes will vary across populations, suggesting that selection against male-beneficial *cis-*regulatory alleles of *Md-HEATR2* on the proto-Y may be population-specific.

The expression divergence of the proto-X and proto-Y copies of *Md-HEATR2* could constitute an early stage of X-Y differentiation before chromosome-wide X-Y differentiation occurs (Bachtrog 2013). Young Y chromosomes have very similar gene content as their ancestral autosomes. In contrast, old Y chromosomes are more likely to have only retained genes with male-specific functions or recruited genes associated with testis expression from other autosomes (Koerich et al. 2008; Kaiser et al. 2011; Mahajan and Bachtrog 2017). Our results suggest that changes in the expression of individual Y-linked genes that were retained from the ancestral autosome could have important phenotypic effects during early Y chromosome evolution, consistent with gene-by-gene divergence on the young Y chromosomes (Wei and Bachtrog 2019).

### Conclusions

We investigated gene sequence and expression differences between the house fly III^M^ proto-Y chromosome and its homologous proto-X to determine how a very young proto-Y/proto-X pair diverge shortly after the proto-Y was formed. To those ends, we used genotypic (III^M^/III) and sex-reversed (III/III) males because they are phenotypically almost the same and only differ in whether they carry a proto-Y chromosome (Hediger et al. 2010; Son et al. 2019). We observe elevated heterozygosity on the proto-sex chromosome in genotypic males (Figure 1), which could be indicative of chromosome-wide sequence differentiation between the proto-Y and proto-X. Alternatively, elevated heterozygosity in genotypic males could merely be a result of being heterozygous for the third chromosome. Consistent with this alternative hypothesis, when we previously substituted a third chromosome without *Mdmd* onto a common genetic background, the effect on gene expression was comparable to substituting a III^M^ chromosome on the same background (Son et al. 2019).

Our subsequent analyses suggest that the house fly III^M^ proto-Y chromosome is differentiated in sequence and expression from its homologous proto-X chromosome at individual genes, but not chromosome-wide. This is consistent with previous work that found evidence for gene-by-gene divergence in a young neo-Y chromosome (Wei and Bachtrog 2019). For example, we only identified two genomic loci (containing a total of 7 genes) that are sufficiently differentiated to assemble into separate proto-X and proto-Y contigs (Figure 2). Notably, 5/7 of those genes have mitochondrial functions, suggesting that mitochondrial function in sperm might be an important target of male-specific selection during the early evolution of a Y chromosome. We did not identify any fixed differences in the protein-coding sequences between the proto-X and proto-Y gametologs of the five mitochondrial genes (Supplementary Table 3), or in the sequence of *Md-HEATR2* (Figure 4), providing further evidence for minimal differentiation between the proto-X and proto-Y. This also suggests that sex-specific selection may be operating on *cis* regulatory sequences of these genes. Consistent with this hypothesis, two of the mitochondrial genes are differentially expressed between the proto-Y and proto-X (Supplementary Table 2). Despite these intriguing examples, there is only a moderate excess of genes with evidence for differential expression between the proto-Y and proto-X (Figure 3). The number of genes with ASE on the third chromosome only in genotypic males, and not in sex-reversed males, is small; we identify fewer than 100 genes that meet these criteria, which is less than 5% of the 2,524 genes assigned to the third chromosome (Meisel and Scott 2018).

We identified one gene on the third chromosome (*Md-HEATR2*) with ASE in genotypic males that is non-ASE in sex-reversed males and has discordant sex-biased expression between genotypic and sex-reversed males (Figure 4). We hypothesize that expression divergence of *Md-HEATR2* could be an example of very early X-Y differentiation of individual genes that results from sex-specific selection. Notably, *Md-HEATR2* is expected to have important functions in spermatogenesis or sperm motility (Diggle et al. 2014; Fuller 1993), similar to the five mitochondrial genes contained within genomic regions that are divergent between the proto-Y and proto-X. Male fertility is an important target of selection during the evolution of old Y chromosomes (Carvalho et al. 2009; Hughes et al. 2010), and our results suggest that selection on male fertility is also an important driver of X-Y divergence during the early evolution of sex chromosomes. Our results also suggest that these selection pressures during the earliest stages of Y chromosome evolution drive gene-by-gene, rather than chromosome-scale, changes in gene expression.

## Materials and Methods

### Fly strain

We analyzed RNA-seq data and performed Oxford Nanopore sequencing on a house fly strain that allows for identification of genotypic (III^M^/III) males and sex-reversed (III/III) males (Hediger et al. 2010). This is because the standard third chromosome (III) in this strain has the recessive mutations *pointed wing* and *brown body*. Sex-reversed males (and normal females) have both mutant phenotypes, whereas genotypic males are wild-type for both phenotypes because the III^M^ chromosome has the dominant wild-type alleles. The RNA-seq data that we analyzed (available at NCBI GEO accession GSE126689) comes from a previous study that used double-stranded RNA (dsRNA) targeting *Md-tra* to create sex-reversed phenotypic males that have a female genotype without a proto-Y chromosome (Son et al. 2019). This is because active *Md-tra* drives female development and inactive *Md-tra* triggers male development (Hediger et al. 2010). We compared gene expression in the abdomens of sex-reversed males, genotypic males that received a sham treatment of dsRNA targeting GFP, and genotypic females that are phenotypically female. We analyzed data from three replicates of each genotype and treatment, with each replicate consisting of a single fly abdomen (Son et al. 2019). We used genotypic males and sex-reversed males from the same strain that were subjected to the same treatment for genome sequencing using the Oxford Nanopore long read technology.

### Oxford Nanopore sequencing

We performed Oxford Nanopore sequencing of one genotypic (III^M^/III) male and one sex-reversed (III/III) male created from the same strain and using the same *Md-tra* dsRNA treatment as a previous RNA-seq study (Son et al. 2019). DNA was isolated with a phenol/chloroform protocol (see Supplementary Materials). Oxford Nanopore Sequencing libraries were prepared with the 1D genomic DNA Ligation kit (SQK-LSK109, Oxford Nanopore), following the manufacturer’s protocol. DNA from the genotypic male and sex-reversed male was used to create a separate sequencing library for each genotype. Following the manufacturer’s protocol, 15 uL of each library, along with sequencing buffer and loading beads (totaling 75 uL), were separately loaded onto two different R9.4 flow cells (i.e., the libraries from the genotypic and sex-reversed males were run on separate flow cells) until no pores were available on a MinION sequencer (Oxford Nanopore).

We performed two different base calling pipelines, using the Guppy pipeline software version 3.1.5 (Oxford Nanopore). First, for the IDP-ASE analysis, we used parameter options with “-- calib_detect --qscore_filtering --min_qscore 10”. Second, for the genome assembly we used the default parameters. We used different parameters for IDP-ASE because base quality affects accurate haplotyping used to estimate ASE (see below), and we therefore used a higher threshold for the base quality. For the IDP-ASE analysis, the base called reads were aligned to the house fly genome assembly v2.0.2 (Scott et al. 2014) using Minimap2 version 2.17 with the “-ax map-ont” parameter (Li 2018).

### Genome assembly, transcript alignment, and sequence divergence

We used wtdbg2 to assemble our base called Oxford Nanopore reads using the default parameters (Ruan and Li 2020). To find genic regions in our genome assembly, we aligned house fly transcripts (from Annotation Release 102) as a query against our genome assembly contigs using BLAT with a minimum mapping score of 50 (Kent 2002). We considered BLAT alignments to be gene copies only if the matched sequence length in the BLAT alignments covers at least half of the original (annotated) transcript length. We selected contigs in our assembly that contain genes that are assigned to the third chromosome in the reference genome.

We used two approaches to differentiate the proto-Y and proto-X contigs. In both approaches we specifically focused on pairs of contigs that contain the same genes because they are indicative of X-Y divergence (i.e., one contig likely contains a proto-Y sequence, and the other contains a proto-X sequence). First, we tested if one contig contains a copy of *Mdmd*, which allows us to designate that contig as the proto-Y copy (and the other is proto-X). *Mdmd* is not present in the annotated house fly genome because the genome was sequenced from female DNA (Scott et al. 2014). To find *Mdmd* copies in our genome assembly, we used the sequence of the *Mdmd* transcript (Sharma et al. 2017) as a query to find contigs containing *Mdmd* with BLAST using an e-value cutoff of 1e-3 (Altschul et al. 1990). Second, we used a *k*-mer comparison approach to differentiate the proto-X and proto-Y contigs (Carvalho and Clark 2013). In this approach, we used a *k*-mer size of 15 to measure the percent unmatched by female reads (%UFR) for every contig in our assembly, with higher values indicating an increased likelihood that a contig is Y-linked. The female reads were from the genome project (BioProject accession PRJNA176013; Scott et al. 2014), and we included the III^M^ male Oxford Nanopore sequencing reads we generated for validation of the bit-array. We followed the options suggested to identify Y sequences in *Drosophila* genomes (Carvalho and Clark 2013), as described previously (Meisel et al. 2017). We used the R package Gviz (Hahne and Ivanek 2016) to visualize genes that are present in III^M^ (proto-Y) contigs and III (proto-X) contigs.

### Variant calling

We used available RNA-seq data (Son et al. 2019) to identify genetic variants (SNPs and small indels) that differentiate the III^M^ proto-Y chromosome from the standard third (proto-X) chromosome, and then we tested if III^M^ males have elevated heterozygosity on the third chromosome as compared to sex-reversed males (Meisel et al. 2017). We used the Genome Analysis Toolkit (GATK) pipeline for calling variants in the RNA-seq data from the *Md-tra* RNAi experiment in (Son et al. 2019), following the best practices for SNP and indel calling on RNA-seq data (McKenna et al. 2010; Meisel et al. 2017). First, we used STAR (Dobin et al. 2013) to align reads from three genotypic (III^M^/III) male libraries and three sex-reversed (III/III) male libraries to the (female) reference assembly v2.0.2 (Scott et al. 2014). The reference genome was sequenced from the aabys strain, which differs from the one we used in our experiment and has males that carry the Y^M^ proto-Y chromosome. The aligned reads were used to generate a new reference genome index from the detected splice junctions in the first alignment run, and then a second alignment was performed with the new reference. We next marked duplicate reads from the same RNA molecule and used the GATK tool ‘SplitNCigarReads’ to reassign mapping qualities to 60 with the ‘ReassignOneMappingQuality’ read filter for alignments with a mapping quality of 255. Indels were detected and realigned with ‘RealignerTargetCreator’ and ‘IndelRealigner’. The realigned reads were used for base recalibration with ‘BaseRecalibrator’ and ‘PrintReads’. The base recalibration was performed in three sequential iterations in which recalibrated and filtered reads were used to train the next round of base recalibration, at which point there were no beneficial effects of additional base recalibration as verified by ‘AnalyzeCovariates’. We next used the recalibrated reads from all three replicates of genotypic and sex-reversed males to call variants using ‘HaplotypeCaller’ with emission and calling confidence thresholds of 20. We applied ‘genotypeGVCFs’ to the variant calls from the two types of males for joint genotyping, and then we filtered the variants using ‘VariantFiltration’ with a cluster window size of 35 bp, cluster size of 3 SNPs, FS > 20, and QD < 2. The QD filter simultaneously considers read depth and variant quality so that a separate read depth filter is not needed during variant filtration. Because this filtration is applied during the joint genotypic step, all variants are genotyped using the combined information across both types of males, eliminating the need to separately cross-reference read mapping information across genotypes. The final variant calls were used to identify heterozygous variants within genes and to estimate ASE with the IDP-ASE tool (see below) using the coordinates from the genome sequencing project, annotation release 102 (Scott et al. 2014). If a locus has different variants (e.g. a reference allele and an alternative allele), we considered the locus heterozygous. We measured relative heterozygosity within each gene in genotypic (III^M^/III) and sex-reversed (III/III) males as the number of heterozygous variants in genotypic males for a given gene (*hG*) divided by the total number heterozygous variants in both genotypic and sex-reversed males (*hSR*), times one hundred: 100*hG*/(*hG* + *hSR*). To annotate each variant within coding sequences, we used SnpEff with filter options “-onlyProtein”, “-no-intergenic”, “-no-downstream”, “-no-upstream”, “-no-intron”, and “-no-utr” (Cingolani et al. 2012).

For the variant calling from Nanopore long reads, the base called reads were indexed using fast5 files with the ‘index’ module of Nanopolish version 0.11.1 (Quick et al. 2016), and they were aligned with Minimap2 version 2.17 (Li 2018) to house fly genome assembly v2.0.2 (Scott et al. 2014). The aligned and raw reads were used to call variants using the “variants” module of Nanopolish version 0.11.1 with the “--ploidy 2” parameter (Quick et al. 2016). We used a python script ‘nanopolish_makerange.py’ provided in the package to split the genome into 50 kb segments because it was recommended to use the script for large datasets with genome size more than 50 kb.

### Allele-specific expression

ASE is the unequal expression of the maternal and paternal alleles in a diploid. Estimating ASE with a single reference genome generates bias in the ASE measurement because RNA-seq reads from the allele found in the reference genome could preferentially map to the reference genome relative to reads from alternative alleles (Stevenson et al. 2013). This read-mapping bias can be reduced by filtering out clusters of variants found in close proximity (Stevenson et al. 2013; Zimmer et al. 2016). We accomplished this by excluding all clusters of 3 or more SNPs found within windows of 35 bp (see above), which is similar to the recommended filtering parameters to reduce mapping bias (Stevenson et al. 2013). Our filtering step retained only 21.3% of all SNPs, which we consider to be high confidence SNPs for inferring ASE. In comparison, Zimmer et al. (2016) applied a filter of 6 SNPs per 100 bp, which we find retains 30.4% of all SNPs in our data. Therefore, the filter we have applied to remove SNP clusters is as stringent or more stringent than those previously used to reduce mapping biases that could affect measurements of ASE.

We investigated if there is elevated ASE on the third chromosome in males carrying one III^M^ proto-Y and one proto-X chromosome compared to sex-reversed males with two proto-X chromosomes. To do this, we implemented the IDP-ASE tool at the gene level with house fly genome annotation release 102 (Scott et al. 2014), following the developers’ recommended analysis steps (Deonovic et al. 2017). The IDP-ASE software was supplied with raw and aligned reads created by RNA-seq (Son et al. 2019) and Nanopore sequencing, as well as variant calls (SNPs and small indels) in RNA-seq reads created by GATK. We only used the Nanopore reads for phasing haplotypes in the IDP-ASE run, and not for variant calling, because there was less than 10× coverage across the house fly genome (i.e., too low for reliable variant calling).

The prepared data from each gene was next run in an MCMC sampling simulation to estimate the haplotype within each gene with a Metropolis-Hastings sampler (Bansal et al. 2008). The software estimates the proportion of each estimated haplotype that contributes to the total expression of the gene (ρ) from each iteration using slice sampling (Neal et al. 2003). A value of ρ=0.5 indicates equal expression between two alleles, whereas ρ<0.5 or ρ>0.5 indicates ASE. The MCMC sampling was run with a 1000 iteration burn-in followed by at least 500 iterations where data were recorded. The actual number of iterations was automatically adjusted by the software during the simulation to produce the best simulation output for quantifying ASE within a gene. The IDP-ASE simulation generated a distribution of ρ for each gene across all post-burn-in iterations, and then it calculated the proportion of iterations with ρ > 0.5. This proportion was used to estimate the extent of ASE for each gene. For example, if all iterations for a gene have ρ > 0.5, then the proportion is 1 and the gene has strong evidence for ASE of one allele. Similarly, if all iterations for a gene have ρ < 0.5, then the proportion is 0 and the gene has strong evidence for ASE of the other allele. In contrast, if half of the iterations have ρ > 0.5 and the other half have ρ < 0.5, then the proportion is 0.5 and there is not any evidence for ASE. In our subsequent analysis, we only included genes with at least 10 mapped reads combined across three RNA-seq libraries from the genotype under consideration.

We used the output of IDP-ASE to compare expression of the III^M^ (proto-Y) and III (proto-X) alleles in genotypic males. IDP-ASE only quantifies ASE within bi-allelic loci, so we only included genes with heterozygous sites within transcripts in genotypic (III^M^/III) or sex-reversed (III/III) males. In addition, we removed heterozygous variants with the same genotype in genotypic and sex-reversed males because they do not allow us discriminate between the proto-Y and proto-X alleles. Removing these variants may have also sped up the simulation times, but this was not rigorously investigated. To discriminate between the III^M^ and III alleles, we used haplotypes estimated during IDP-ASE runs and genotypes inferred from GATK for genotypic (III^M^/III) and sex-reversed (III/III) males. For example, using genotypes called using GATK from the RNA-seq data, we first identified sites with heterozygous alleles in genotypic males and homozygous alleles in sex-reversed males. Next, we inferred the allele in common between genotypic and sex-reversed as the III allele, and the other allele that is unique to genotypic males as the III^M^ allele. Lastly, we matched those sites to the haplotypes estimated by IDP-ASE to quantify ASE within each genotype.

## Acknowledgements and funding information

We thank Leo W. Beukeboom and Daniel Bopp for collaborating on the RNAi knockdown and valuable discussions, and Fei Yuan for help developing a python script used to calculate the percentage of heterozygous variants. This work was supported by a Grant-in-Aid of Research from the National Academy of Sciences, administered by Sigma Xi, The Scientific Research Society (grant number G2018100198487895 to J.H.S. and R.P.M.) and by the National Science Foundation (grant numbers OISE-1444220 and DEB-1845686 to R.P.M.). Analysis of RNA-seq and Oxford Nanopore sequencing data were performed on the Maxwell and Sabine clusters in the Research Computing Data Core at the University of Houston. The Oxford Nanopore long read data used in the study are available from the National Center for Biotechnology Information Sequence Read Archive under BioProject accession PRJNA620357 (BioSample accessions SAMN14518459 for sex-reversed males and SAMN14518460 for genotypic males). The III^M^ male genome assembly is available from the National Center for Biotechnology Information Whole Genome Shotgun accession JACCFA000000000 under BioProject accession PRJNA620357.

## Supplementary Materials

### Identification of two candidate proto-X/proto-Y loci

We identified one contig containing *Mdmd* in the III^M^ assembly (ctg2382) that contains the same three genes as another contig (ctg1607) without *Mdmd* (Figure 2A). We infer the contig with *Mdmd* to be the proto-Y sequence, and the one without to be from the proto-X. There are two additional contigs containing *Mdmd* in our III^M^ genome assembly, but we did not find any corresponding proto-X contigs for these (Supplementary Figure 1). These other two contigs with *Mdmd* either contain loci without sufficient X-Y divergence to assemble into separate proto-X and proto-Y contigs, or we did not sequence deep enough to capture enough proto-X reads to assemble the proto-X gametolog. The male-determining regions of the house fly proto-Y chromosomes contain both complete and truncated copies of *Mdmd* (Sharma et al. 2017), and ctg2382 contains two truncated copies of *Mdmd*. The genomic region corresponding to this contig is found in a single scaffold of the reference genome (NW_004774683.1), which was previously assigned to the third chromosome (Meisel and Scott 2018). There are three genes at this locus, and they are present in the same order in the reference genome, the proto-Y contig, and proto-X contig (Figure 2A).

We found the second locus using a *k*-mer comparison approach to test for contigs containing sequences that are enriched in males relative to females (Carvalho and Clark 2013). Here, we compared *k-*mers in our Nanopore III^M^ male reads and the Illumina reads used to generate the reference (female) genome assembly (Scott et al. 2014) to our III^M^ male assembly. We then calculated the percent of each contig in our III^M^ assembly that is unmatched by female reads (%UFR), which we expect to be 100% for a Y chromosome contig that only contains male-specific *k-*mers. We did not identify any contigs with greater than 27.4%UFR (Supplementary Figure 1), which is consistent with the minimal sequence divergence between the proto-Y and proto-X chromosomes (Meisel et al. 2017). To determine an expectation for what constitutes a %UFR of a proto-Y contig in our assembly, we calculated %UFR for the three contigs that each contain copies of *Mdmd* (Supplementary Figure 1). The largest %UFR of any contig with *Mdmd* is 13.8% (ctg3705). We used this value as a threshold to find proto-Y contigs that do not contain *Mdmd*.

We found 275 contigs with a %UFR>13.8, nine of which (Supplementary Table 1) contain protein-coding genes that were previously assigned to the third chromosome (Meisel and Scott 2018). One gene (LOC101898200, which encodes histone H3) was found on six different contigs with %UFR>13.8 (and an additional 27 contigs with %UFR<13.8). Histone gene clusters are a common feature of most metazoan genomes, and the multiple copies are under tight regulatory control (McKay et al. 2015; Duronio and Marzluff 2017).These contigs with high %UFR and copies of histone H3 genes are either false positives or not of substantial phenotypic relevance. Two of the other contigs with %UFR>13.8 (ctg3539 and ctg8407) each contain 1-2 genes that are found as multiple copies on other contigs (all with a %UFR<13.8). One of those genes (LOC109612838 on ctg3539) encodes a protein with predicted reverse transcriptase and RNase H activity, suggesting that it is a retrotransposon (Finnegan 2012). Another gene (LOC109613819 on ctg8407) encodes a protein with zinc finger and integrase domains, also consistent with a transposable element (Volff 2006). We exclude these contigs from our subsequent analysis because they likely reflect transposable element expansions within the house fly genome. We only found one contig with a %UFR>13.8 (ctg2522, which has 16.9%UFR) that has a single corresponding contig (ctg1519) with the same four genes (Supplementary Table 1). We assigned ctg2522 to the proto-Y chromosome and ctg1519 to the proto-X (Figure 2B). All four genes shared by both contigs are mapped to the third chromosome in the reference genome. These four genes are present in the same order and same orientation in the reference genome (on scaffold NW_004764689.1), the proto-Y contig, and proto-X contig.

### DNA isolation with phenol/chloroform protocol

A single genotypic male and a single sex-reversed male with detached wings were each transferred to a 1.5 mL tube with 0.5 mL homogenization buffer (4.1 g sucrose, 15 mL 1M Tris-HCl pH 8.0, 0.5M EDTA, 100 mL dH2O) and then homogenized using a pestle set into a tissue grinder homogenizer. To each tube we added 40 uL of 10% SDS and 2.5 uL of 10 mg/mL Proteinase K, and then we incubated the tube at 65°C for 30 min. We next added 2 uL of 4mg/mL RNase to each tube and incubated at 37°C for 15 min. We added 48 uL of 5M KAc to each tube and placed on ice for 30 min. Then we centrifugated the tubes at 14000 rpm for 10 min at 4°C, and the supernatant was transferred into a new tube using a wide-bore pipette tip (all subsequent steps of transfer and mixing during DNA extraction were also done with wide-bore pipette tips to prevent DNA shearing). We added 250 uL phenol and 250 uL chloroform to the extracted supernatant in the new tube, mixed briefly, spun at 14000 rpm for 15 min at 4°C, and then transferred the supernatant into a new tube. We next added 500 uL chloroform to the supernatant in the new tube, mixed well, and spun a 14000 rpm for 5 min at 4°C. We then transferred the supernatant into a new tube. We added 40 uL of 3M NaAc and 800 uL of 95% ethanol to the supernatant in the new tube, mixed briefly, spun at 14000 rpm for 15 min at 4°C, and then carefully poured off all supernatant. We next added 800 uL of 70% ethanol to the remaining pellet, mixed briefly to wash the pellet, spun at 14000 rpm for 15 min at 4°C, removed the supernatant, and then resuspended the pellet in 30 uL of nuclease-free water.

**Supplementary Figure 1.**
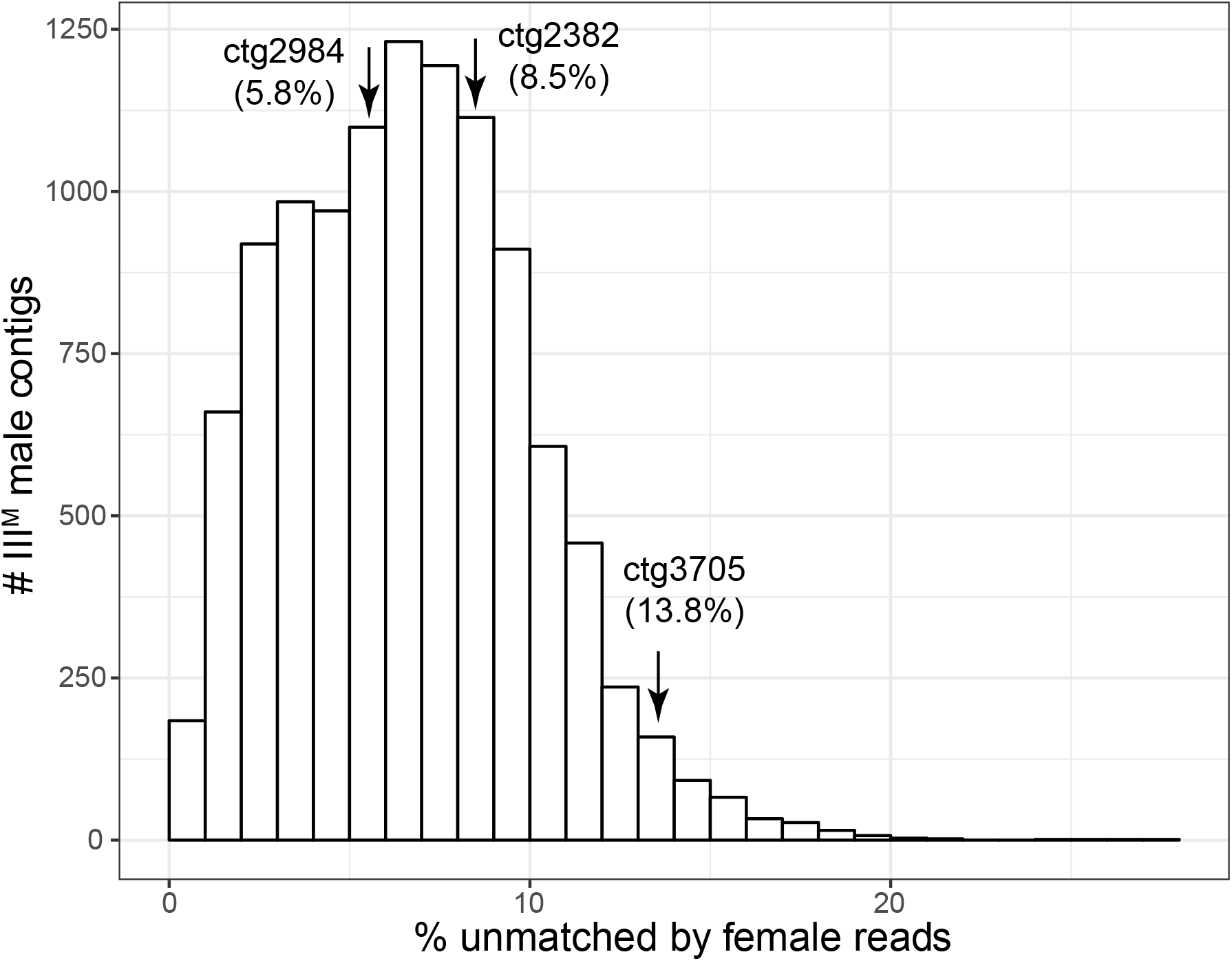
Histograms of the percent unmatched by female reads (%UFR) for all contigs in the III^M^ male genome assembly. Three contigs with *Mdmd* are shown, with their %UFR in parentheses.

**Supplementary Figure 2.**
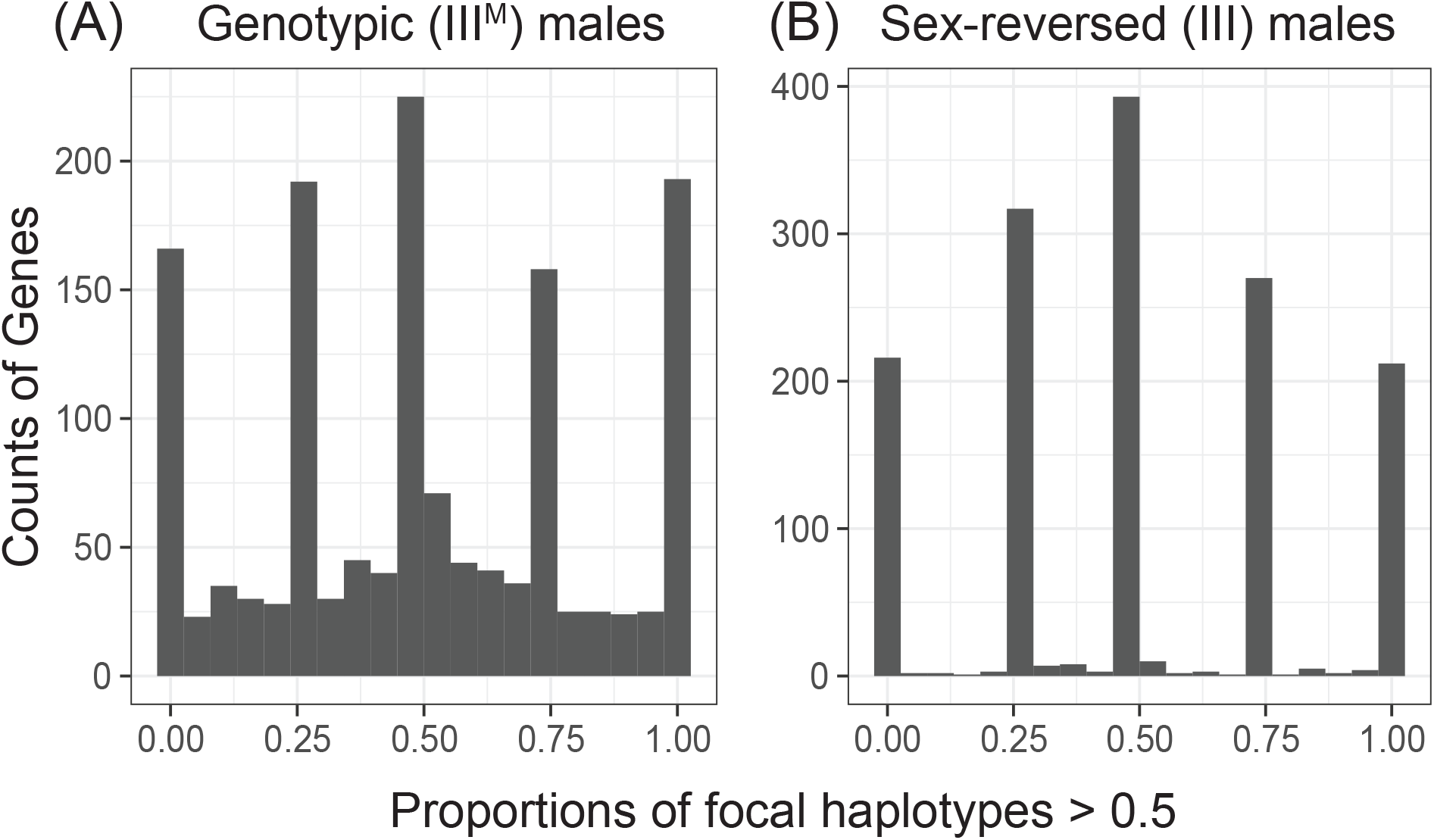
Histograms of third chromosome genes in different ASE categories. ASE was measured in (A) genotypic (III^M^/III) and (B) sex-reversed (III/III) males. ASE is measured as the proportion of iterations in which a focal haplotype is >0.5 of the alleles expressed across iterations in an MCMC simulation. We consider a gene to have ASE if the proportion is <0.125 or >0.875. Genes with a proportion of focal haplotypes between 0.375 and 0.625 are classified as not having ASE.

**Supplementary Figure 3.**
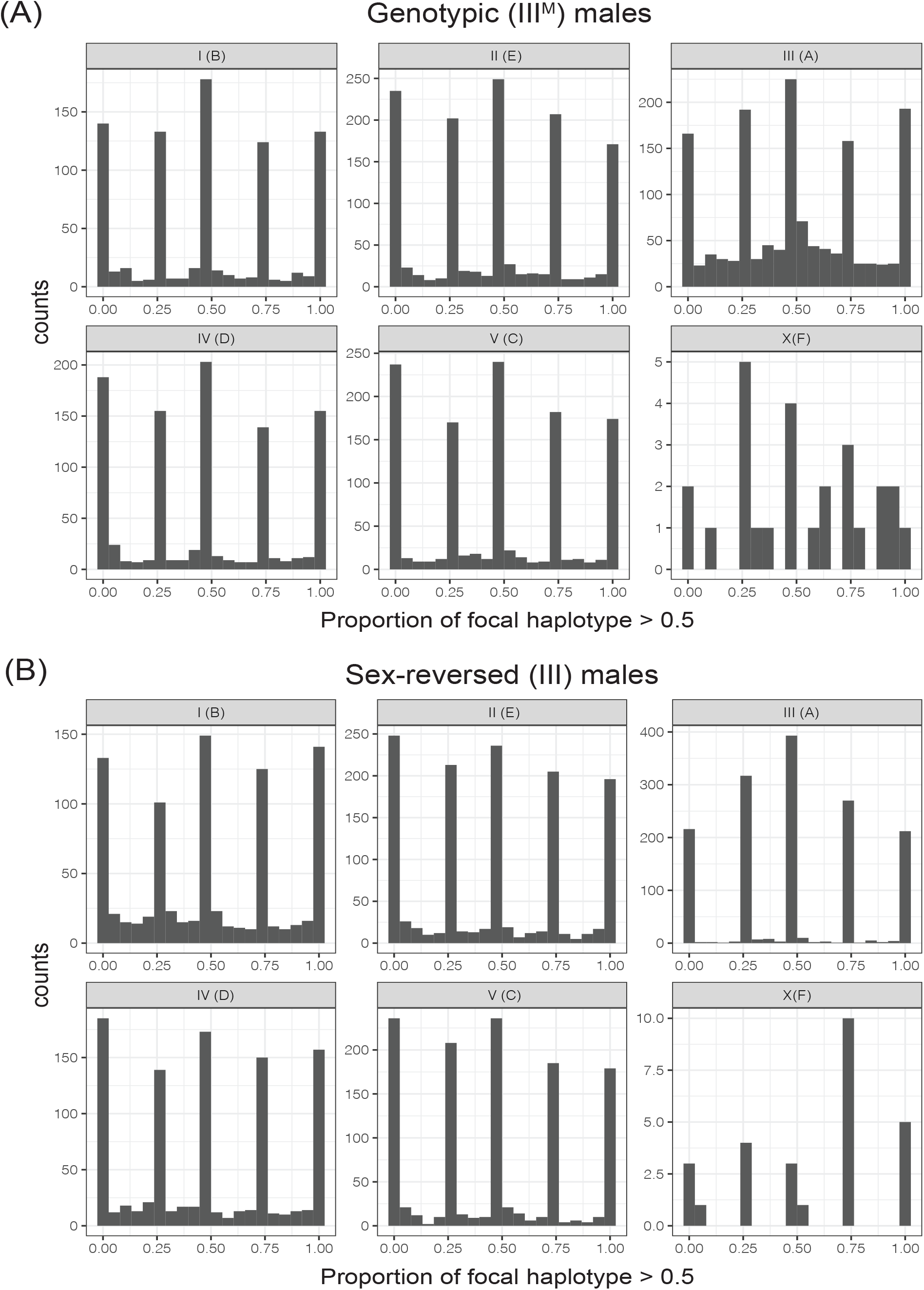
Histograms of allele-specific expression (ASE) in genes for all chromosome are shown in the genotypic (III^M^/III) males and the sex-reversed (III/III) males. Muller element nomenclature (from *Drosophila*) for each chromosome is given in parentheses (Meisel and Scott 2018). If a gene is expressed equally between two alleles, the proportion of the focal haplotype is 0.5; otherwise, the proportion is greater or less than 0.5.

**Supplementary Table 1.** Nine contigs in the III^M^ male assembly contain genes that are assigned to the third chromosome and have a %UFR>13.8. The genes on each contig are listed.

| Contig ID | Gene | # copies of gene | %UFR of contig |
| --- | --- | --- | --- |
| ctg2522 | LOC101894374 | 1 | 16.9 |
|  | LOC101894024 | 1 |  |
|  | LOC101894537 | 1 |  |
|  | LOC101894698 | 1 |  |
| ctg3539 | LOC101888429 | 1 | 15.4 |
|  | LOC109612838 | 1 |  |
| ctg8407 | LOC109613819 | 1 | 16.6 |
| ctg1423 | LOC101898200 | 6 | 14.5 |
| ctg4658 | LOC101898200 | 2 | 14.0 |
| ctg5526 | LOC101898200 | 1 | 17.3 |
| ctg8144 | LOC101898200 | 1 | 14.8 |
| ctg9654 | LOC101898200 | 1 | 20.1 |
| ctg10432 | LOC101898200 | 1 | 15.9 |

**Supplementary Table 2.**
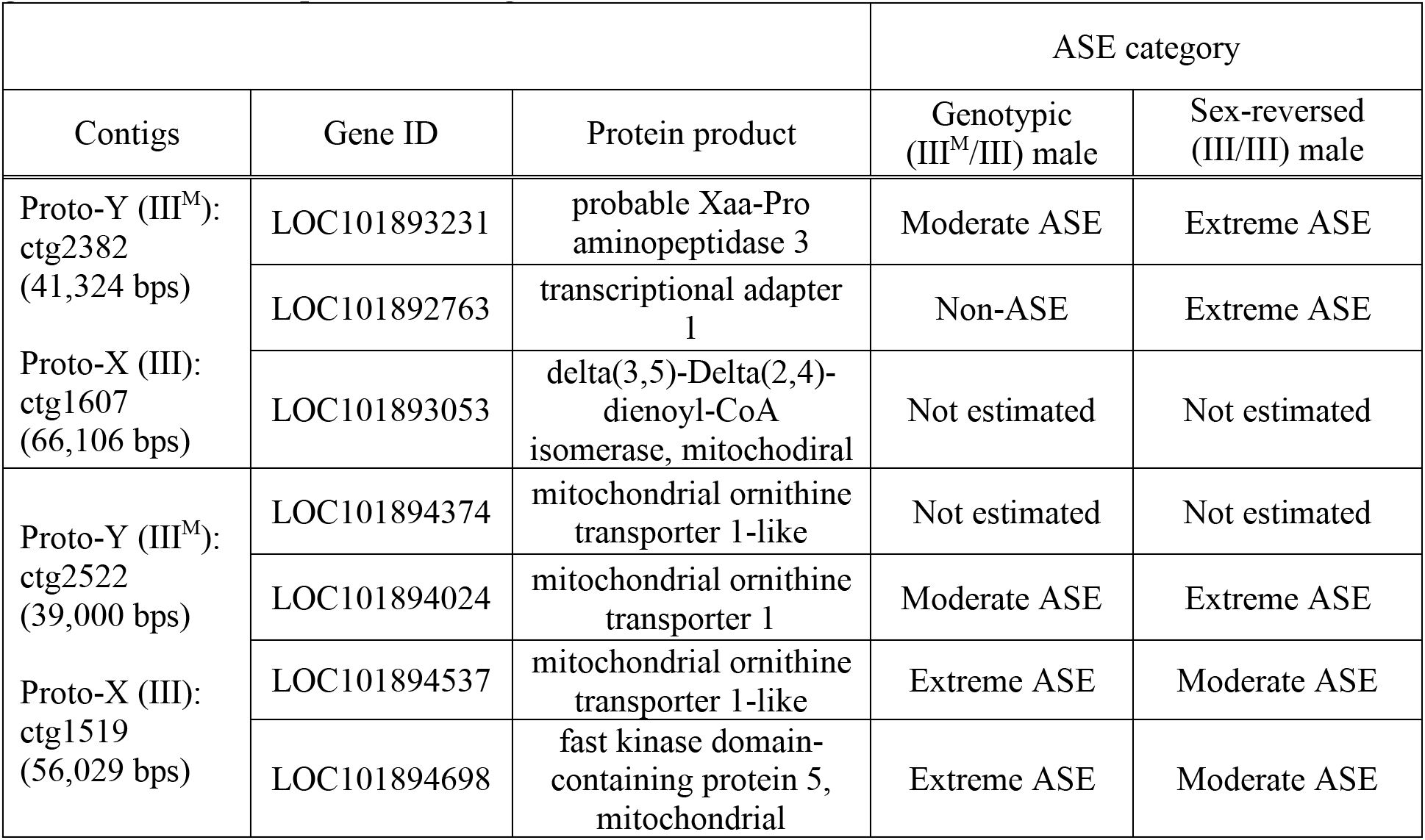
Genes on contigs with differentiated proto-Y (III^M^) and proto-X (III) copies. There are two pairs of contigs (each pair contains a proto-Y contig and a proto-X contig). Results of a test for ASE in genotypic (III^M^/III) males are shown in the second to last column and ASE of the sex-reversed (III^M^/III) males in the last column. ASE was not estimated for some genes due to low sequence coverage.

**Supplementary Table 3.** Missense alleles in coding sequences of two genes on the proto-Y and proto-X contigs. LOC101894698 has also five variable sites that are all synonymous, which are not shown in the table. Variants in coding sequences of the other three genes on the proto-Y and proto-X (LOC101892763, LOC101894024, and LOC101894537) are synonymous and not shown in the table.

| Gene | Position at scaffold | III<sup>M</sup> male genotype | III male genotype | III<sup>M</sup> allele | Reference genome allele |
| --- | --- | --- | --- | --- | --- |
| LOC101894698 | 117354 | T/A | A/A | T | T |
|  | 118649 | C/T | T/T | C | C |
|  | 118909 | T/C | C/C | T | T |
| LOC101893231 | 17382 | C/T | C/C | T | C |
|  | 17429 | A/G | A/A | G | A |
|  | 29021 | G/A | A/A | G | G |

**Supplementary Table 4.** Counts of ASE genes and non-ASE genes on each chromosome in genotypic (III^M^/III) males and sex-reversed (III/III) males.

| Chromosome<br>(Muller<br>element) | # genes with<br>ASE in<br>III<sup>M</sup> males | # genes with<br>non-ASE in<br>III<sup>M</sup> males | # genes with<br>ASE in<br>III males | # genes with<br>non-ASE in<br>III males | Odds<br>ratio | 95% CI of<br>odds ratio |
| --- | --- | --- | --- | --- | --- | --- |
| III(A) | 456 | 413 | 438 | 412 | 1.038574 | 0.8555646<br>-<br>1.2606663 |
| **genome**<br>**except**<br>**III(A)** | **1635** | **1089** | **1711** | **1010** | **0.886281** | **0.7933539**<br>-<br>**0.9900421** |
| I(B) | 320 | 222 | 334 | 212 | 0.915002 | 0.7123784<br>-<br>1.1750187 |
| II(E) | 465 | 314 | 513 | 291 | 0.840134 | 0.6821523<br>-<br>1.0344483 |
| IV(D) | 394 | 249 | 395 | 215 | 0.861373 | 0.6798614<br>-<br>1.0908786 |
| V(C) | 448 | 296 | 460 | 288 | 0.947652 | 0.7654588<br>-<br>1.1730543 |
| X(F) | 8 | 8 | 9 | 4 | 0.720539 | 0.102255<br>-<br>4.754933 |

**Supplementary Table 5.** Counts of genes with ASE on each chromosome in genotypic (III^M^/III) males and sex-reversed (III/III) males. The total number of genes (# genes) in each chromosome group with ASE measurements in both genotypic and sex-reversed males (second column), # genes with ASE in genotypic males and non-ASE in sex-reversed males (third column), and # genes with non-ASE in genotypic males and ASE in sex-reversed males (fourth column) are shown. Bold indicates statistical significance (*P* < 0.05)

|  |  | ASE in III<sup>M</sup> males<br>and non-ASE in<br>III males | non-ASE in<br>III<sup>M</sup> males and<br>ASE in III males | Fisher's Exact Test |  |
| --- | --- | --- | --- | --- | --- |
| Chromosome<br>(Muller<br>element) | #<br>genes | # genes | # genes | Odds ratio<br>compared<br>with III(A) | 95% CI |
| III(A) | 1420 | 95 | 76 |  |  |
| genome<br>except III(A) | 4201 | **241** | **281** | **1.45665** | **1.014895 -**<br>**2.095777** |
| I(B) | 824 | 51 | 59 | 1.444139 | 0.8688985 -<br>2.4078676 |
| II(E) | 1236 | **68** | **86** | **1.578592** | **0.9961489 -**<br>**2.5100267** |
| IV(D) | 966 | **55** | **76** | **1.724145** | **1.063163 -**<br>**2.809137** |
| V(C) | 1149 | 67 | 77 | 1.434875 | 0.8982881 -<br>2.2981210 |
| X(F) | 26 | 0 | 0 | 0 | 0 - Infinity |

**Supplementary Table 6.**
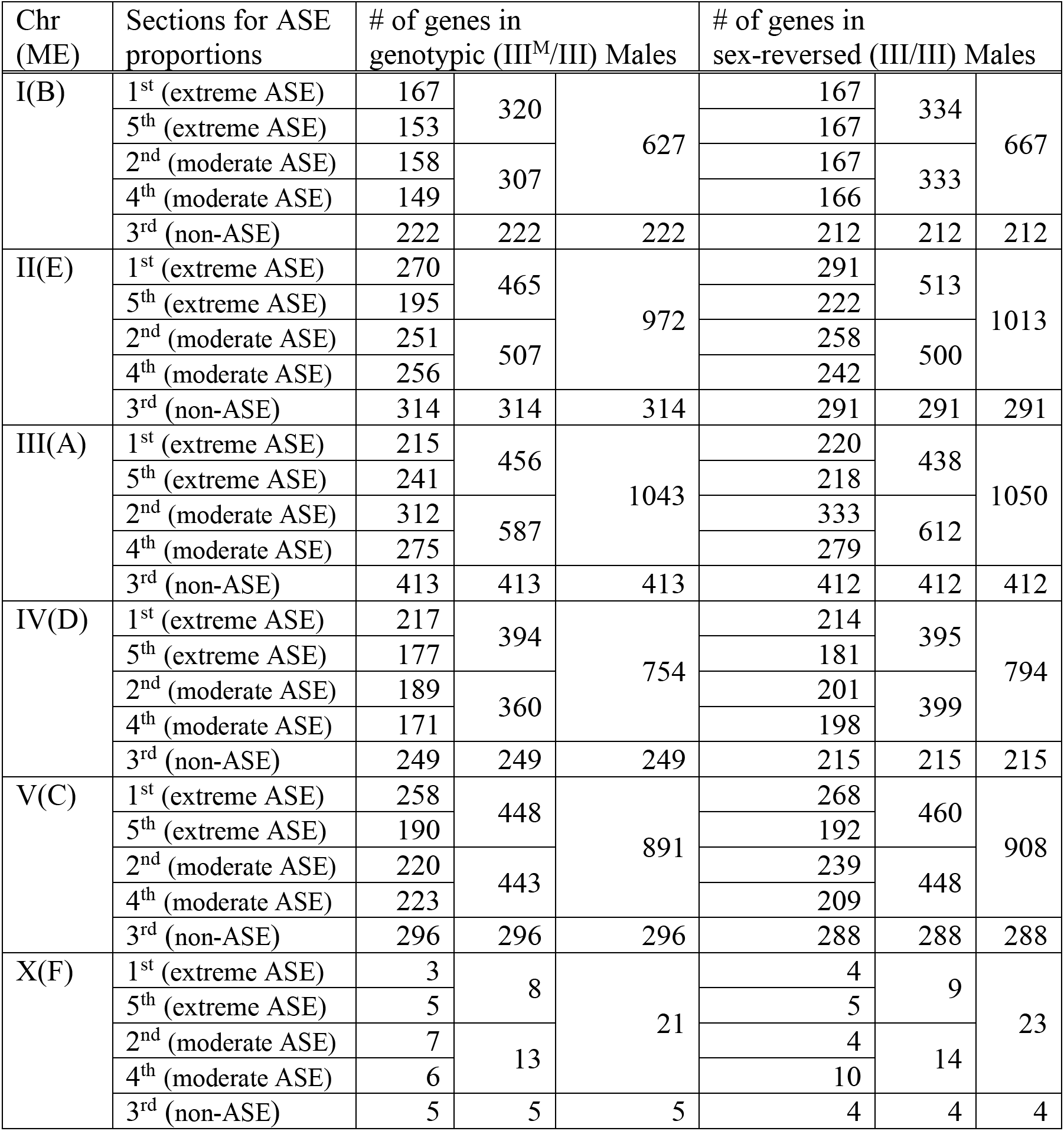
Counts of ASE genes based on the division of ASE measurements into five bins, following the rules described in the Methods. ASE proportions are sorted in the order of extreme (1^st^ and 5^th^), moderate (2^nd^ and 4^th^), and no (3^rd^) ASE. Only extreme ASE was used in the comparisons with non-ASE presented in the main text.

**Supplementary Table 7.** The number of annotated genes assigned to each chromosome (Muller elements in parentheses) and total length of the corresponding chromosomes in base pairs (bp).

| Chromosomes<br>(Muller elements) | # genes on<br>chromosomes | Size in length (bp) |
| --- | --- | --- |
| I(B) | 2128 | 73015049 |
| II(E) | 3175 | 95424831 |
| III(A) | 2271 | 76855929 |
| IV(D) | 2400 | 76690558 |
| V(C) | 2660 | 81074832 |
| X(F) | 47 | 1486543 |
| Unassigned | 3859 | 345856202 |
| Total | 16540 | 750403944 |

**Supplementary Data 1.** A VCF file called with the RNA-seq reads

**Supplementary Data 2.** A VCF file called with the Oxford Nanopore reads

**Supplementary Data 3.** k-mer comparison in the III^M^ male genome assembly

**Supplementary Data 4.** IDP-ASE (allele-specific expression) output for the genotypic males

**Supplementary Data 5.** IDP-ASE (allele-specific expression) output for the sex-reversed males

Supplementary Data 1 and 2 are available at Dryad (https://doi.org/10.5061/dryad.280gb5mnk or https://datadryad.org/stash/share/6f4xn0dRAu8j2SQXCmqKICv95gYfpGzlhoR5SYLJZy0)

